# Interleukin-1β and -6 release from immune cells by DOPA-based melanin as free pigment or complexed to carboxymethylcellulose

**DOI:** 10.1101/185140

**Authors:** Koen P. Vercruysse, Tonie S. Farris, Margaret M. Whalen

**Affiliations:** Chemistry Department, Tennessee State University, Nashville, TN 37209

**Keywords:** DOPA, interleukin, melanin, polysaccharide

## Abstract

We have observed that many polysaccharides can promote the oxidation of 3,4-dihydroxyphenylalanine (DOPA) into melanin-like pigments leading to the formation of water-soluble polysaccharide/melanin complexes. These pigments were characterized by size exclusion chromatography and FT-IR spectroscopy. The effect on the secretion of interleukin (IL)-lβ and IL-6 from immune cells by DOPA-based melanin synthesized in the presence or absence of carboxymethylcellulose (CMC) was evaluated. We observed that the melanin/CMC complex had a more potent effect on both IL secretions compared to the melanin prepared from DOPA in the absence of any polysaccharide. The study of the effect of melanins on the IL secretion by immune or other cells will help illuminate the potential contributions of this broad class of pigments to pathological conditions like Parkinson’s disease or ochronosis.

## 1. Introduction

Melanins (MNs) are darkly colored pigments ubiquitously found in nature and excellent reviews regarding their biosynthesis, chemistry, classification and functions have been written.[1] The precise chemical structure of the MNs is not firmly established. Based upon degradation reactions involving oxidation in an alkaline environment, various model structures have been proposed and it has been suggested that MNs should be described as “heterogenous polymers derived by the oxidation of phenols and subsequent polymerization of intermediate phenols and their resulting quinones”.[1] MNs are broadly divided into eumelanin and pheomelanin. The eumelanins typically found in animals are built from L-DOPA, an oxidation product of L-tyrosine and have a dark brown to black color. Pheomelanins are built from a combination of DOPA and L-cysteine and are typically reddish in color.[1] The biosynthesis of eumelanin from L-DOPA involves a sequence of oxidation, cyclization and polymerization reactions most commonly described by the Raper-Mason scheme. [2] A similar process has been discussed for dopamine or norepinephrine, and these catecholamines (CAs) together with L-DOPA may be the precursors of the so-called neuromelanins.[1, 3, 4] Neuromelanins are considered an aberration from the normal metabolization pathways for these CAs and have been associated with oxidative stress conditions. [5-7] In addition to L-DOPA or CAs, other catechols can serve as precursors for different types of MNs. MNs are also produced by plants, but the precursors of these pigments are nitrogen free phenols like catechol, caffeic acid, chlorogenic acid or others. Such plant pigments are responsible, e.g., for the browning observed when plant or fruit materials are cut. [8] Apart from animals and plants, MN-like pigments are also produced by fungi or bacteria.[1] Homogentisic acid, a para-diphenol rather than an ortho-diphenol, is the precursor for the dark pigment observed in the later stages of alkaptonuria. The pathology associated with this pigment deposition is termed ochronosis.[9]

Using DOPA or CAs as precursors, MNs can readily be synthesized in an alkaline environment in the presence of an oxidizing agent.[10-12] However, such synthetic MNs are often insoluble and difficult to characterize or test for biological activities. We have explored the possibility that MNs could be synthesized in the presence of polysaccharides (PS) leading to water-soluble PS/MN complexes, providing more suitable preparations for biological evaluations.[13]

Interleukin 1 beta (IL-1β) is a cytokine that stimulates the inflammatory response as well as cellular growth and tissue reconstruction.[14-17] IL-1β is secreted by a variety of cell types including; monocytes, macrophages, T cells, natural killer (NK) cells, keratinocytes, and fibroblasts.[14, 16, 18, 19] IL-1β can act as a co-stimulator of the production of interferon gamma (IFNγ), another pro-inflammatory cytokine, in natural killer (NK) cells.[20] This multiple-action cytokine contributes to chronic inflammation in diseases such as rheumatoid arthritis and multiple sclerosis[21, 22] Chronic inflammation also contributes to cancer cell growth and elevated IL-1β levels occur in breast and pancreatic cancers and melanoma.[23-27] Poor prognoses in patients with breast, colon, and lung cancers as well as melanomas have been associated with increased IL-1β production.[19, 26] Conversely, IL-1β is an essential component of the response to infection and thus tight regulation of its levels are required for overall health.[17] Thus, any agent that causes dysregulation of IL-1β levels has the potential to cause either unwanted inflammation or an inability to mount a competent immune response to infection. IL-6 is a glycosylated protein that is synthesized as a precursor protein of 212 amino acids.[28] IL-6 is produced mainly by monocytes/macrophages, fibroblasts, and vascular endothelial cells. [29] Other cells that can express this cytokine include keratinocytes, osteoblasts, T cells, B cells, neutrophils, eosinophils, mast cells, smooth muscle cells, and skeletal muscle cells.[29, 30] IL-6 regulates endometrial tissue growth, bone formation, and hormone synthesis.[31, 32] Elevation of IL-6 can be seen in inflammatory diseases such as rheumatoid arthritis, systemic lupus erythematosus, psoriasis, and Crohn’s disease. [33] High IL-6 levels have been shown in patients with carcinomas such as breast, lung, and lymphoma[34]; as a result, studies have linked elevation of IL-6 levels with tumor progression.[35, 36]

Using synthetic MNs[37] and MNs isolated from human brains[38]or herbal sources[39, 40], many studies have shown that (neuro)melanins may act as proinflammatory agents. Therefore we undertook a study of the effect of a DOPA-based MN on the interleukin release from immune cells. One such type of DOPA-based MN was synthesized in the presence of NaOH and yielded a soluble eumelanin. Earlier experiments performed in our laboratories indicated that many polysaccharides (PS) can promote the formation of MN from CAs.[13] Thus, in analogy with our experiments involving CAs, a second type of DOPA-based MN was prepared through an air-mediated oxidation of DOPA in the presence of carboxymethylcellulose (CMC) without the addition of any base. CMC was chosen as this is a synthetic, pharmaceutical grade PS that is readily available and inert. The synthesis of MNs in the presence of PS leads to highly water-soluble PS/MN complexes, with the MN possibly covalently attached to the PS backbone. Both types of DOPA-derived MNs were characterized by size exclusion chromatography (SEC) and FT-IR spectroscopy and tested for their capacity to affect IL-1β or IL-6 release from immune cells.

## 2. Materials and Methods

### 2.1 Materials

Carboxymethylcellulose (low viscosity grade; sodium salt), fucoidan (from *Fucus vesiculosus)* and D,L-3,4- dihydroxyphenylalanine (DOPA) were obtained from Sigma-Aldrich (Milwaukee, WI). L-ascorbic acid was obtained from Fisher Scientific (Suwanee, GA). All other reagents were of analytical grade.

### 2.2 Synthesis and purification of MNs #1 and #2

MN #1 was prepared in a 5mL mixture containing 16mM DOPA and 0.15M NaOH, kept at room temperature for thirty days. MN #2 was prepared by adding 216mg DOPA to a 500mL solution containing 10 mg/mL CMC. This mixture was left stirring at 37°C for two days. Both MNs were dialyzed using Spectrum Spectra/Por RC dialysis membranes with molecular-weight-cut-off of 3.5kDa obtained from Fisher Scientific (Suwanee, GA, USA) against water (up to 3.5L) for three days with up to four changes of water each day. The dialyzed mixtures were kept at -20°C overnight and dried for three days using a Labconco FreeZone Plus 4.5L benchtop freeze-dry system obtained from Fisher Scientific (Suwanee, GA, USA).

### 2.3 UV_Vis spectroscopy

UV/Vis spectroscopic measurements were made in wells of a 96-well microplate using the SynergyHT microplate reader from Biotek (Winooski, VT). All measurements were performed at room temperature (RT) using 200 μL aliquots and 200μL water as the blank.

### 2.4 Size exclusion chromatography (SEC)

SEC analyses were performed on a Breeze 2 HPLC system equipped with two 1500 series HPLC pumps and a model 2998 Photodiode array detector from Waters, Co (Milford, MA, USA). Analyses were performed using an Ultrahydrogel 500 column (300 X 7.8 mm) obtained from Waters, Co (Milford, MA, USA) in isocratic fashion using a mixture of 25mM Na acetate:methanol:acetic acid (90:10:0.05% v/v) as solvent. Analyses were performed at room temperature and 20 μL of sample was injected.

### 2.5 FT-IR spectroscopy

FT-IR spectroscopic scans were made using the NicoletiS10 instrument equipped with the SmartiTR Basic accessory from ThermoScientific (Waltham, MA). Scans were taken with a resolution of 4 cm^-1^ between 650 and 4,000 cm^-1^ at room temperature using a KBr beam splitter and DTGS KBr detector. Each spectrum represents the accumulation of 32 scans.

### 2.6 Preparation of monocyte-depleted (MD) peripheral blood mononuclear cells (PBMCs)

PBMCs were isolated from Leukocyte filters (PALL-RCPL or RC2D) obtained from the Red Cross Blood Bank Facility (Nashville, TN) as described elsewhere. [41] Leukocytes were retrieved from the filters by back-flushing them with an elution medium (sterile PBS containing 5 mM disodium EDTA and 2.5% [w/v] sucrose) and collecting the eluent. The eluent was layered onto Ficoll-Hypaque (1.077g/mL) and centrifuged at 1,200g for 50 min. Granulocytes and red cells pelleted at the bottom of the tube while the PBMCs floated on the Ficoll-Hypaque. Mononuclear cells were collected and washed with PBS (500g, 10min). Following washing, the cells were layered on bovine calf serum for platelet removal. The cells were then suspended in RPMI-1640 complete medium which consisted of RPMI-1640 supplemented with 10% heat-inactivated BCS, 2 mM Z-glutamine and 50 U penicillin G with 50 μg streptomycin/mL. This preparation constituted PBMCs. Monocyte-depleted PBMCs (10-20% CD16+, 10-20 % CD56+, 70-80% CD3+, 3-5% CD19+, 2-20% CD14+) were prepared by incubating the cells in glass Petri dishes (150 × 15 mm) at 37 °C and air/CO_2_, 19:1 for 1 h. This cell preparation is referred to as MD-PBMCs.

### 2.7 PBMC-pigment studies

Test compounds were dissolved in water one day prior to the start of the experiment. For both IL-1β and IL-6 release experiments, 25 μL of the test compounds (and the appropriate controls) were added to 500 μL of MD-PBMC cell suspension. The concentration of the cells was 750,000 in 500 μL of cell culture media. After addition of the compounds, the mixtures were incubated at 37°C in an atmosphere containing 5% CO_2_ for 24 h after which cell viability was assessed and supernatants were collected and stored at -80°C until assay.

### 2.8 Cell Viability

Cell viability was assessed at the end of each exposure period. Viability was determined using the trypan blue exclusion method. Briefly, cells were mixed with trypan blue and counted using a hemocytometer. The total number of cells and the total number of live cells were determined for both control and treated cells to determine the percent viable cells.

### 2.9 IL-1β or IL-6 release assays

Aliquots from the PMBC cells exposed to pigment were diluted 20-fold for IL-1β analysis or 400-fold for IL-6 analysis. IL-1β and IL-6 levels in the diluted aliquots were assessed using the OptEIA™ enzyme-linked immunosorbent assay (ELISA) for human IL-1β kit and human IL-6 respectively (BD Biosciences, San Diego, CA). In brief, capture antibody diluted in coating buffer was applied to wells of 96-well flat-bottom micro-plates specifically designed for ELISA (Fisher Scientific). The plates were incubated overnight at 4°C, and then excess capture antibody was removed by washing the plate three times with wash buffer (PBS containing 0.05% [v/v] Tween-20 (PBST). The wells were then treated with blocking buffer to prevent non-specific binding and the plate was sealed and incubated at room temperature for 1 hr. Blocking buffer was removed with three washes of PBST, and cell supernatants and IL-1β or IL-6 standards were added to dedicated wells; the plate was re-sealed and incubated at room temperature for 2 hr. The plate was then washed five times with PBST and this was followed by incubation with IL-1β or IL-6 detection antibody which was subsequently conjugated with horseradish peroxidase. The plate was then washed seven times and a substrate solution was added for 30 min at room temperature to produce a colored product. The incubation with the substrate was ended by addition of acid and the absorbance was measured at 450 nm on a Thermo Labsystems Multiskan MCC/340 plate reader (Fisher Scientific).

## 3. Results

### 3.1 Preliminary observations

Preliminary observations made in our laboratory indicated that DOPA can be converted into MN-like pigments in the presence of PS, similar to what had been established for CAs. Without the addition of base or redox-sensitive ions like Cu^2+^, DOPA undergoes very little auto-oxidation to MN. However, in the presence of select PS, a bright orange colored solution is generated in the initial phase of the reaction, followed by a gradual darkening. The formation of MN from DOPA in the presence of select PS is illustrated in Fig. 1. Reaction mixtures were prepared in wells of a cell culture dish by dissolving about 30mg DOPA in 10mL water and adding about 100mg PS or other chemicals. The mixtures were left at room temperature and Fig. 1 shows a photograph taken after three days of reaction.

**Fig. 1:**
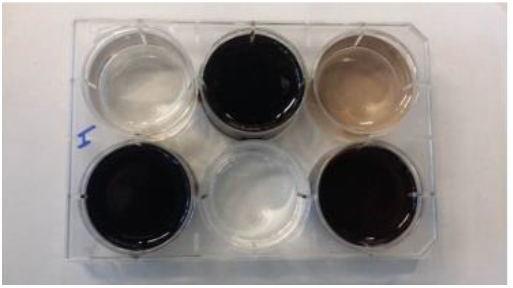
Photograph of reaction mixtures containing about 30mg DOPA, dissolved in 10mL water, and about 100mg test substance after three days at room temperature. Test substances were: top, left to right, none, sodium acetate and sodium glucuronate; bottom, left to right, fucoidan, L-ascorbic acid and CMC.

Black solutions or gels (due to high viscosity) were obtained in the presence of sodium acetate, fucoidan and CMC. A light brown solution was obtained in the presence of sodium glucuronate, and a colorless solution was obtained in the presence of L-ascorbic acid. In the case of sodium acetate, a significant amount of pigment was observed to be adhered to the bottom of the well after siphoning off the dark solution. This was not observed in the pigment solutions prepared in the presence of fucoidan or CMC. The soluble portions of these reaction mixtures could be dialyzed against water and lyophilized to a dry powder for further characterization. In addition to fucoidan and CMC, other PS, e.g., chondroitin sulfate or alginate, were capable of generating water-soluble PS/pigment complexes using DOPA as the precursor (results not shown).

### 3.2 Synthesis of MNs #1 and #2

MN #1 and #2 were prepared, as described in Materials and Methods, separately from the experiments illustrated in Fig. 1. The reactions were performed in a glass test tube or flask to minimize the potential adhesion of MN to the sides of the reaction containers. The reactions between DOPA and NaOH (MN #1) or between DOPA and CMC (MN #2) resulted in dark brown to black solutions. SEC analyses were performed to monitor the progress of the chemical reactions and the dialysis purification processes. Fig. 2 presents the SEC profiles, shown at 275 and 400nm, of MNs #1 and #2 following dialysis of the reaction mixtures.

**Fig. 2:**
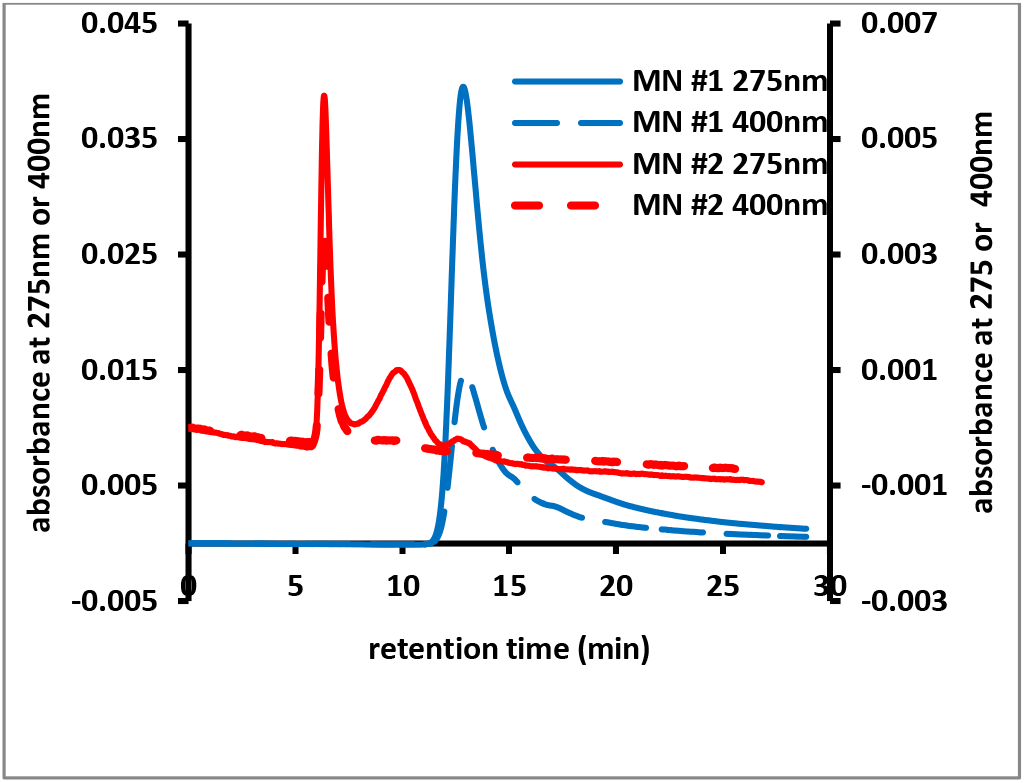
SEC profiles of MN #1 (blue lines and left Y-axis) and MN #2 (red lines and right Y-axis), generated as described in Materials and Methods. The solid lines represent the signal at 275nm, while the dashed lines represent the signal at 400nm.

When lyophilized, MN #1 yielded a black, fine powder and MN #2 yielded a grey-black, fibrous material. Both dried materials were subjected to FT-IR spectroscopy. Fig. 4 presents an overlay of the FT-IR spectra of DOPA and MN #1. Fig. 5 presents an overlay of the FT-IR spectra of CMC and MN #2.

**Fig. 4:**
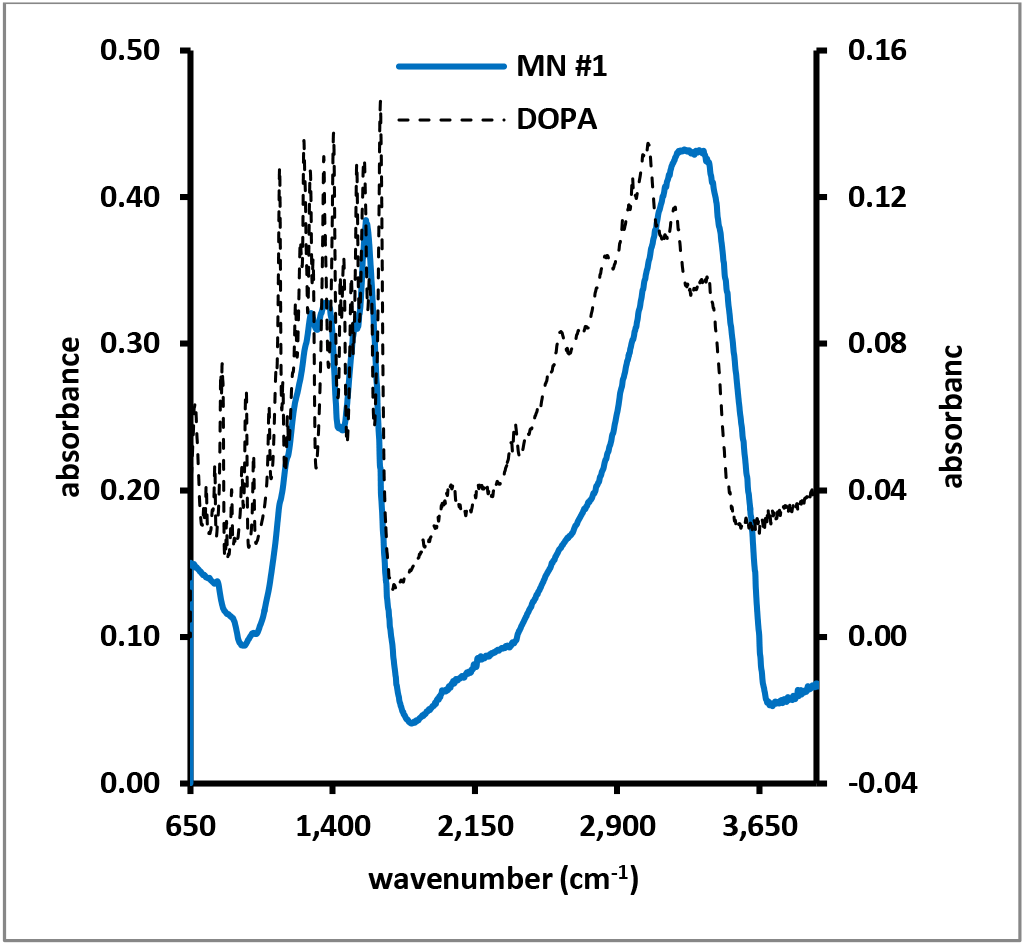
FT-IR spectra of DOPA (dashed light grey line; right Y-axis) and pigment #1 (solid blue line; left Y-axis)

**Fig. 5:**
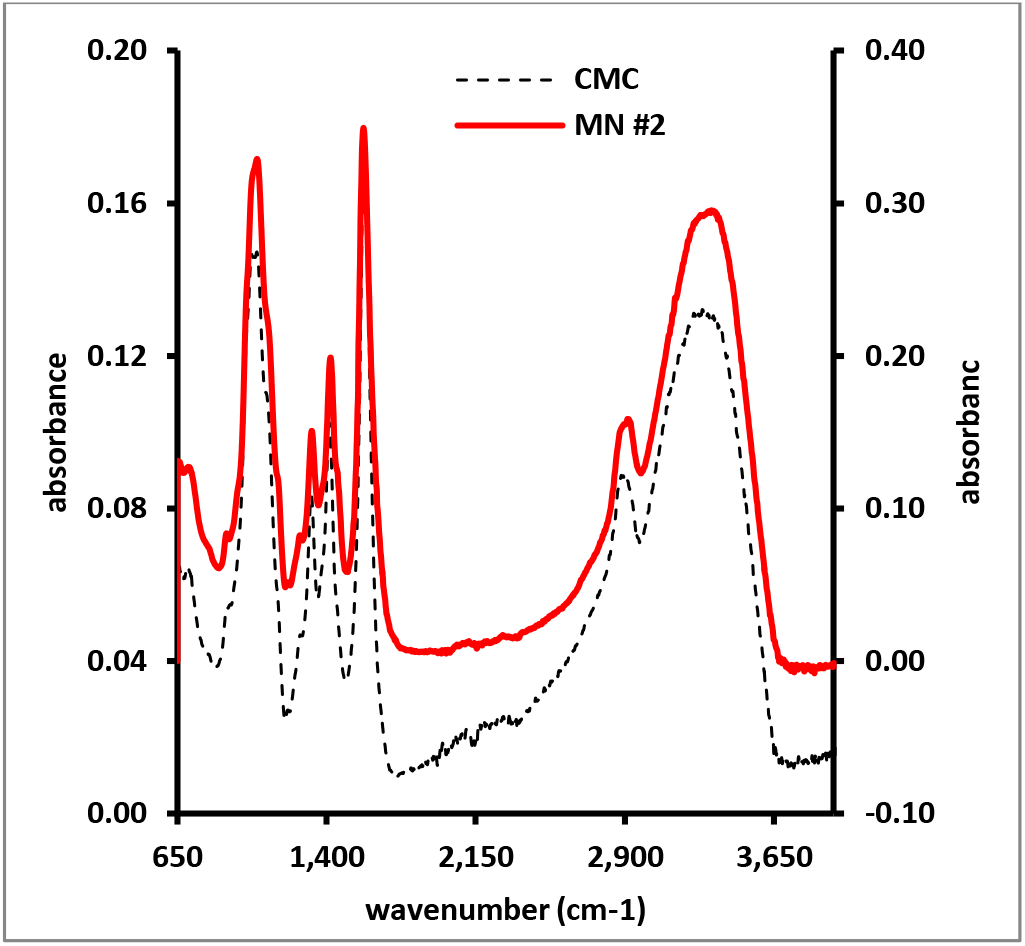
FT-IR spectra of CMC (dashed grey line; right Y-axis) and MN #2 (solid red line; left Y-axis)

In our SEC analyses, the lower limit of exclusion, as determined by the injection of water, was about 15 minutes. Peaks with a retention time lower than 15 minutes are high molecular mass compounds, while peaks with retention times of 15 minutes or higher are low molecular mass compounds. DOPA was determined to have a retention time of about 15 minutes. SEC analyses performed during the reactions revealed, for both reactions, the emergence of a new peak in the high molecular mass range and with absorbance in the visible range of the spectrum. MN #1 had a peak retention time of about 12.75 minutes for the signals at 275nm and 400nm. MN #2 had peak retention times of about 9.45 minutes, visible in the UV range, but not in the Vis range of the spectrum, and a second peak retention time of about 6.3 minutes, visible in both the UV and Vis range of the spectrum. The peak at 9.45 minutes is related to CMC as judged from SEC analyses of CMC standard solutions. The peak at 6.3 minutes represents the DOPA-based MN, possibly complexed to CMC. Fig. 3 presents the UV_Vis spectra of the peaks with absorbance in the Vis range for both MN materials.

**Fig. 3:**
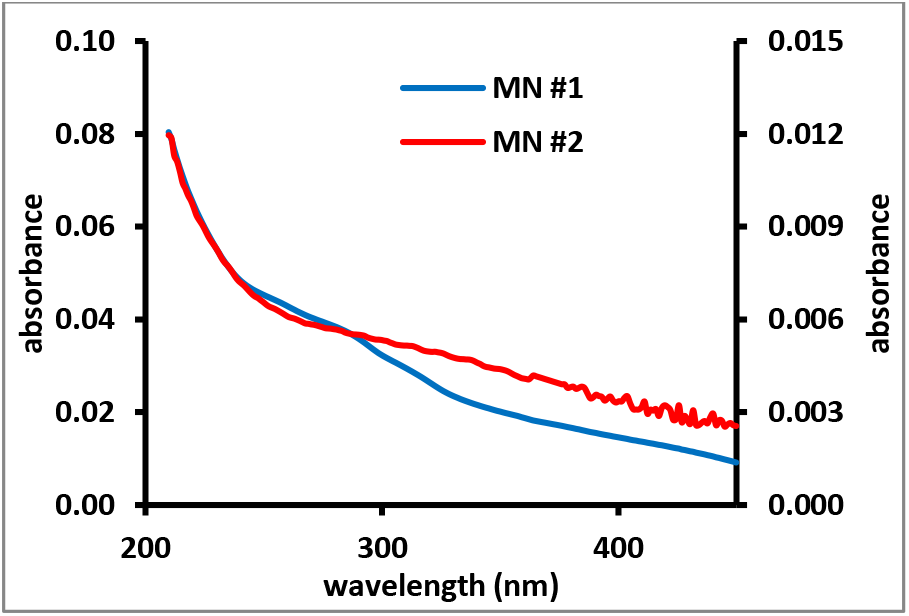
UV_Vis spectra of the peaks in the SEC profiles of MN #1 (blue line; left Y-axis) and MN #2 (red line; right Y-axis) with absorbance in theVis range.

### 3.3 Interleukin release from immune cells

To study the IL release from immune cells, lyophilized MN #1 was dissolved in water and a dilution series with concentrations between 0 and 5 mg/mL was prepared. The absorbance of these dilutions was measured at 400, 425,450, 475 and 500nm and linear relationships (r^2^ > 0.999) between pigment concentration and absorbance were obtained (results not shown). A stock solution of lyophilized MN #2 was prepared at a concentration of 10.4 mg/mL and its absorbance was measured at 400, 425, 450, 475 and 500nm. Using the linear relationships obtained with MN #1 it was estimated that the stock solution of MN #2 contained between 0.14 and 0.25 mg/mL pigment material, thus allowing us to estimate that MN #2 contains between 1.4 and 2.4% pigment. Aliquots of the dilution series made from MN #1 and of the stock solution made from MN #2 were incubated with immune cells and the releases of IL-1β and IL-6 from these immune cells were measured. No cytotoxicity due to the presence of MN #1 or #2 was observed. Fig. 6 illustrates the concentration of IL-1β secreted by the immune cell mixture as a function of the pigment concentration present in the cell mixture.

**Fig. 6:**
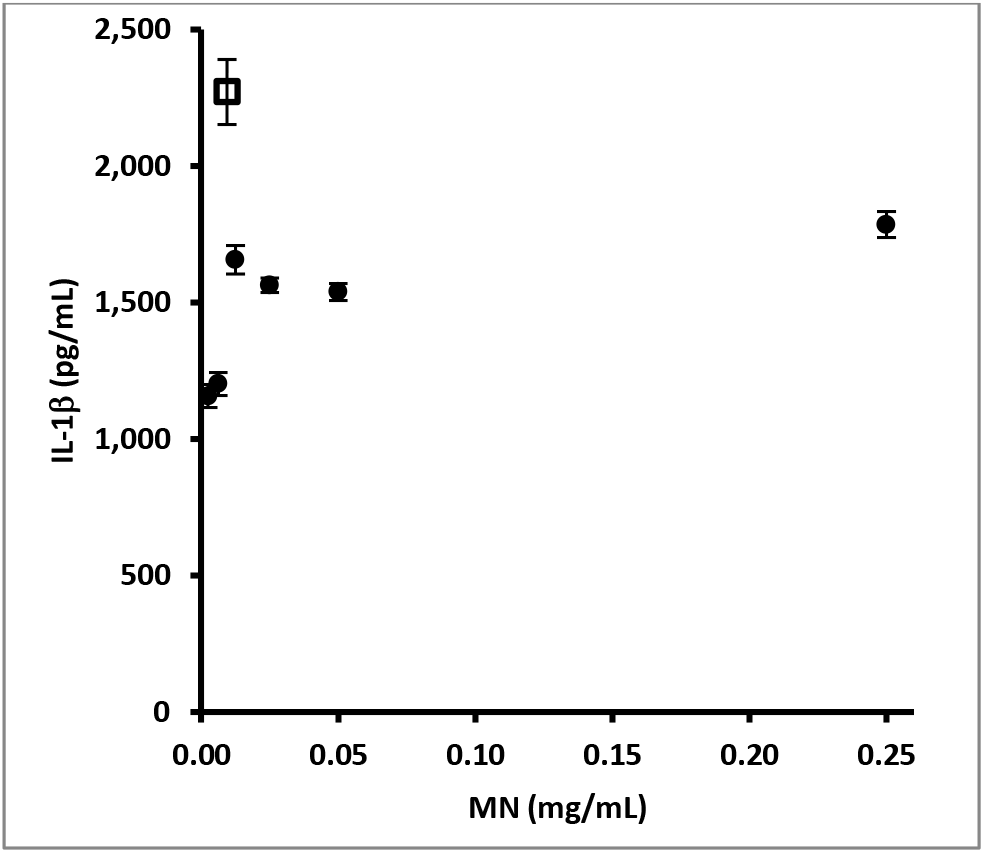
Average (n=3; error bars represent standard deviations) IL-1β concentration secreted by immune cell mixture as a function of MN concentration by MN #1 (black circles) and MN #2 (open square)

Fig. 7 illustrates the concentration of IL-6 present in the immune cell mixture as a function of the pigment concentration present in the cell mixture.

**Fig. 7:**
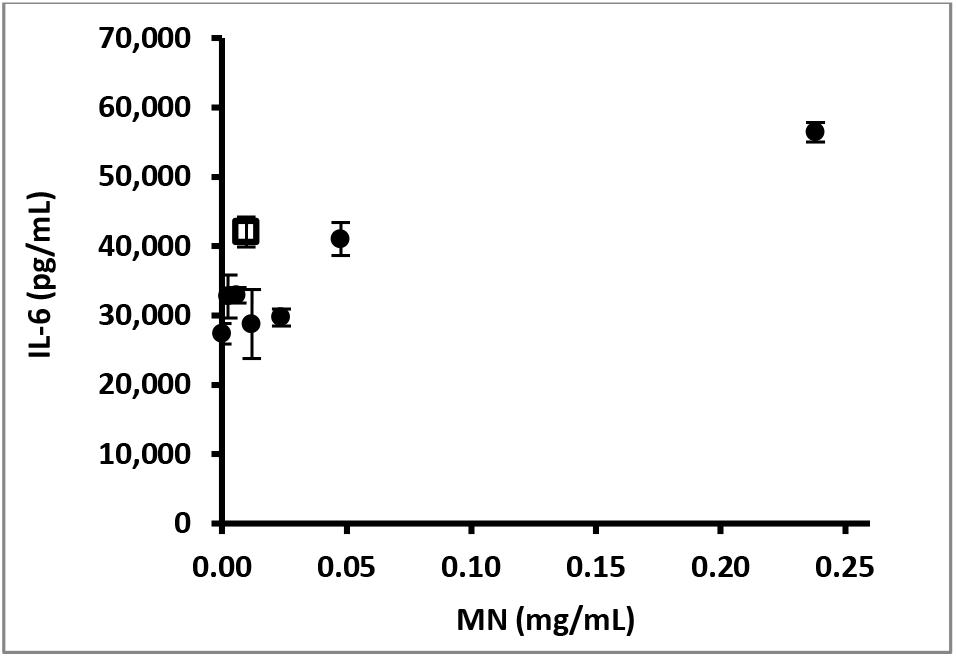
Average (n=3; error bars represent standard deviations) IL-6 concentration secreted by immune cell mixture as a function of pigment concentration by MN #1 (black circles) and MN #2 (open square)

## 4. Discussion

Overall, our observations suggest that PS like CMC and others can promote the oxidation of DOPA to MN-like pigment. Similar observations have been made for other catecholic precursors and the results of these experiments will be presented elsewhere. Although at this time the MN-like materials made in the presence of PS cannot be fully characterized, we have reasons to believe that covalent bonding may exist between the PS and the MN. When MN is synthesized in the absence of PS in plastic containers, then a significant amount of pigment is bound to the surface as discussed for the results shown in Fig. 1. No such sticking to plastic surfaces is observed when the MN is synthesized in the presence of PS. In addition, when solutions of CMC/MN complex are dialyzed against water containing 0.15M NaCl or are extracted with organic solvent like ethylacetate, no pigment is removed from the CMC/MN-containing solution. Finally, dried CMC/MN material readily (within minutes) dissolves in water at a concentration of 10 mg/mL and the final solution has little noticeable viscosity. CMC solutions in water at 10 mg/mL typically require multiple hours to dissolve in to a homogenous, highly viscous solution. The possibility that the MN is covalently attached to the PS backbone is in accordance with other reports that have suggested that MN-like pigments are associated with PS fractions in some organisms, e.g., the binding of MN-like pigments to chitin in the exoskeleton of insects. [42-44]

SEC separates molecules based upon differences in their hydrodynamic volume, often related to the molecular mass of the molecule. Thus, using SEC equipped with PDA detection one can readily make a distinction between low molecular mass compounds, including low molecular mass chromophores, and high molecular mass compounds, including pigments or pigment/PS complexes.[13] The retention time at 6.3 minutes of the peak with absorbance in the Vis range present in the SEC analysis of MN #2 (see Fig. 2) is much lower than the retention time of CMC. This suggests that the hydrodynamic volume of this complex is much higher than that of the original CMC material. This could be due to aggregation of multiple molecules of PS/MN complex or due to the possibility that the PS/MN complex consist of PS-stabilized MN nanoparticles. The possibility that PS-stabilized MN nanoparticles are generated in the reactions described in this report is supported by the fact that other studies have shown that naturally-occurring, MN types of pigments do exist as nanoparticulate material.[45, 46]

The UV_Vis spectra (see Fig. 3) of MN #1 and #2 show similar patterns: an exponential decline of the absorbance profile with strong absorbance in the UV range and weak absorbance in the Vis range. These spectra are in accordance with spectra of MN discussed elsewhere.[2] Comparing the FT-IR spectra of DOPA to MN #1 (see Fig. 4) one observes that the sharp peaks present in the spectrum of DOPA are not present in the spectrum of MN #1. The FT-IR spectrum of MN #1 contains broad peaks between wavenumbers of about 1,200 and 1,600cm^-1^ and a broad peak around wavenumber 3,300 cm^-1^. This pattern is similar to the FT-IR spectra of other DOPA-based MN or the FT-IR spectrum of MN obtained from sepia ink. [47, 48] No qualitative differences can be observed between the FT-IR spectra of CMC and MN #2 (see Fig. 5), despite the fact that MN #2 is grey-black in color. These results suggest that, despite the dark color of the material, very little MN is present and that the bulk of MN #2 consists of CMC. The estimate that MN #2 contains between 1.4 and 2.4% (w/w) pigment, as discussed in the results section, would be in accordance with the observations regarding the comparison of the FT-IR spectra of CMC and MN #2. In addition, similar observations had been made for the synthesis of MN-like materials from CAs in the presence of PS.[13]

The IL-1β release from immune cells as a function of MN #1 concentration appeared to follow a hyperbolic pattern (see Fig. 6). MN #2 appeared to strongly enhance the IL-1 β release from immune cells compared to MN #1. CMC by itself had been determined not to affect the IL-1 β release from immune cells. The IL-6 release from immune cells as a function of MN #1 concentration appears to follow a linear pattern (see Fig. 7). MN #2 appeared to enhance the IL-6 release from immune cells. CMC had been observed not to affect the IL-6 release from immune cells. Thus, MN #2 appeared to affect the release of IL-1β and -6 from immune cells more potently than CMC or MN #1.

In general, our observations suggest that select PS may promote the oxidation of DOPA and its subsequent polymerization to MN-like pigments. Enzymes like tyrosine hydroxylase are often implicated in the formation of MNs, but in some instances there appears to be no correlation between the activity of such enzymes and the density of the MNs observed in tissues. [49] However, it has been observed before that the oxidation of CAs and related compounds is enhanced at higher pH values. [50] This has been confirmed in this work by the formation of (insoluble) DOPA-based pigment in the presence of sodium acetate (see Fig. 1) and the synthesis of (soluble) DOPA-based pigment, MN #1, in the presence of 0.15M NaOH. With regards to the PS-mediated synthesis of MN-like pigments from DOPA, we envision a possible dual role for the PS materials. First, although the pH values of the PS solutions are close to neutral, the PS backbone may provide an alkaline microenvironment, particularly in the case of polyanionic PS like CMC or fucoidan, to promote the oxidation of DOPA. Therefore we hypothesize that anionic PS could act as base catalysts for the oxidation and further polymerization of DOPA or other precursors into MN-like pigments. Secondly, the PS backbone may provide a template to direct the polymerization of oxidized precursors, similar to the, hotly debated, “dirigent model” of the biosynthesis of lignins in plants.[51, 52] Apart from DOPA, we have used PS to synthesize, water-soluble, MN-like pigments from a wide variety of precursors and these results will be presented elsewhere.

Thus, generating water-soluble MN materials using a variety of precursors will help in the study of the cell-biological effects of these pigments, e.g., the IL release from cells as discussed in this report. Apart from being used for external coloration and camouflage, UV protection, etc., MNs are present internally and have been linked to the pathophysiology of Parkinson’s disease, Alzheimer’s disease or ochronosis.[53, 54]

## Acknowledgements

This research was in part supported by NIH grant U54CA163066.

